# Stepwise Ligand Capture Primes PKR for Activation and RNA Discrimination

**DOI:** 10.1101/2025.07.15.665015

**Authors:** Chong Han, Chen Li, Yi-Feng Xu, Ming Rao, Ling-Ling Chen, Jiaquan Liu

## Abstract

Protein Kinase R (PKR) triggers the innate immune response upon detecting double-stranded RNAs (dsRNAs). Its kinase activity is controlled by two dsRNA-binding domains (dsRBDs), yet how these dsRBDs operate during RNA ligand recognition and downstream signaling remains unresolved. Here, we uncover an unexpected interplay between the dsRBDs that orchestrates a two-step ligand sampling, ensuring accurate and efficient PKR activation. dsRBD1 surveys all RNA duplexes through one-dimensional diffusion but dissociates within milliseconds to minimize background signaling, while dsRBD2-RNA binding plays a key role in fostering kinase-kinase interactions. Upon encountering foreign dsRNAs, dsRBD2 arrests the diffusive dsRBD1, thereby establishing a scaffold essential for kinase dimerization and phosphorylation. By contrast, prevalent bulges and large internal loops in self-RNA duplexes obstruct dsRBD2 from capturing dsRBD1, creating a steeply elevated energy landscape that enables PKR to discriminate foreign RNAs from abundant self-RNAs, thus evolving a delicate strategy to achieve both high selectivity and robust signaling.

## INTRODUCTION

RNA-binding proteins (RBPs) are important regulators of various cellular processes. To date, over 2,000 RBPs have been identified, many of which feature RNA-binding domains (RBDs) such as RNA recognition motifs (RRMs), K homology (KH) domains, zinc fingers (ZnFs), and double-stranded RNA-binding domains (dsRBDs)^1,2^. While individual RBDs typically bind short RNA sequence motifs with low affinity^3–5^, tandem RBDs can synergistically enhance binding strength and improve substrate selectivity—often by orders of magnitude^1,6–8^. Such cooperation between RBDs is believed to be essential for the functions of RBPs, yet the molecular basis behind this process is not fully understood^1,6^.

Among those RBPs with tandem RBDs, PKR is a latent serine/threonine kinase that mainly detects and responds to viral infections^9^. PKR consists of three independently folding domains: two N-terminal dsRBDs (dsRBD1 and dsRBD2), and a C-terminal kinase domain (KD) linked to the dsRBDs by a flexible linker. Upon recognizing dsRNA, a common byproduct of viral RNA replication, PKR’s KD undergoes autophosphorylation, leading to its activation^10–12^. Once activated, PKR phosphorylates eukaryotic translation initiation factor 2 alpha (eIF2α), halting global protein synthesis to inhibit viral replication^13^. Beyond its role in antiviral defense, PKR can be aberrantly activated by endogenous RNAs in uninfected cells^14^, and has been implicated in a range of pathological conditions, including neurodegenerative diseases^15–17^, lupus^18^, and metabolic disorders like obesity and diabetes^19,20^.

Under normal conditions, PKR phosphorylation must be tightly controlled through its RNA-binding activity. Consistent with this notion, modulation of PKR–RNA interactions—either by dsRNA mimics acting as decoys^18,21,22^ or by regulatory RBPs such as PACT and ADAR1^23–26^— has emerged as a key safeguard against inappropriate PKR activation. Although RNA recognition lies at the heart of these events, how the dsRBDs engage with the dsRNA in complex has remained unclear^25^. Initial studies suggested that dsRBD2 plays a regulatory role in PKR activation, potentially by maintaining an autoinhibited conformation in the absence of RNA^27,28^. Later work revealed that kinase phosphorylation requires proximity-induced KD dimerization or polymerization^10,11^, a process facilitated by the dsRBD-RNA interactions. More recent studies indicated that PKR binds dsRNA in a highly dynamic manner, underscoring the complexity of dsRBD function in PKR^22,25^. Together, these findings suggest that PKR activation is unlikely to follow a simple "lock-and-key" model proposed for other canonical dsRBDs^29^.

Like PKR, all immune systems must identify foreign ligands amid a vast excess of host-derived ligands, prompting the use of strategies that enhance discrimination to ensure accurate recognition. For example, T cell receptors (TCRs) apply an error-correction mechanism known as kinetic proofreading during the recognition of foreign peptides presented by major histocompatibility complexes (pMHC)^30–32^. For PKR, this task is particularly challenging because viral RNAs typically appear at much lower concentrations^33^ and PKR must respond early in host defense^34^. How the tandem arrangement of dsRBDs contributes to sharpening PKR’s RNA discrimination remains unknown.

In this study, we report a previously unrecognized interplay between the two dsRBDs of PKR. The process begins with dsRBD1 engaging with a dsRNA independently and scanning along the duplex in a one-dimensional (1D) manner. This rapid movement produces only transient contacts that are insufficient to support intermolecular KD-KD interactions. On nonself-RNAs, dsRBD2 traps the sliding dsRBD1, stabilizing the complex, and thus facilitating kinase dimerization. In contrast, self-RNAs often contain bulges or internal loops that interfere with the dsRBD interplay. These observations ultimately reveal a unified model in which the functional coupling of dsRBDs licenses both PKR activation and RNA discrimination.

## RESULTS

### dsRBD2 arrests sliding dsRBD1 on dsRNA

We exploited prism-based single-molecule total internal reflection fluorescence (smTIRF) microscopy to visualize PKR binding to a long dsRNA that mimics viral RNA. Single 11.6-kb dsRNA molecules were constructed by *in vitro* transcription, stretched across a passivated custom-made flow cell surface by laminar flow, and linked at both ends via biotin-neutravidin interactions (**Figures 1A, 1B and S1A**; **Table S1**). Human PKR protein was purified and labeled with a Cy3 fluorophore similar to our previous studies^22,35^ (**Figures 1C and S1B**; **Table S2**). Following the injection of Cy3-PKR into the flow cell, PKR binding on 11.6-kb dsRNA was monitored in real-time (**Figure S1C**), and three types of PKR-dsRNA association mechanics were identified (**Figure 1D**). The most frequent interaction, accounting for 58% of total events, involved the static binding of PKR at a random location on the 11.6-kb dsRNA (**Figures 1D and 1E**, Static). The least frequent interaction, representing only 2% of the events, involved PKR molecules freely diffusing along the long dsRNA (**Figures 1D and 1E**, Mobile). The third type, observed in 40% of the events, showed a dynamic transition between the static and mobile binding (**Figures 1D and 1E**, Static + Mobile).

**Figure 1.**
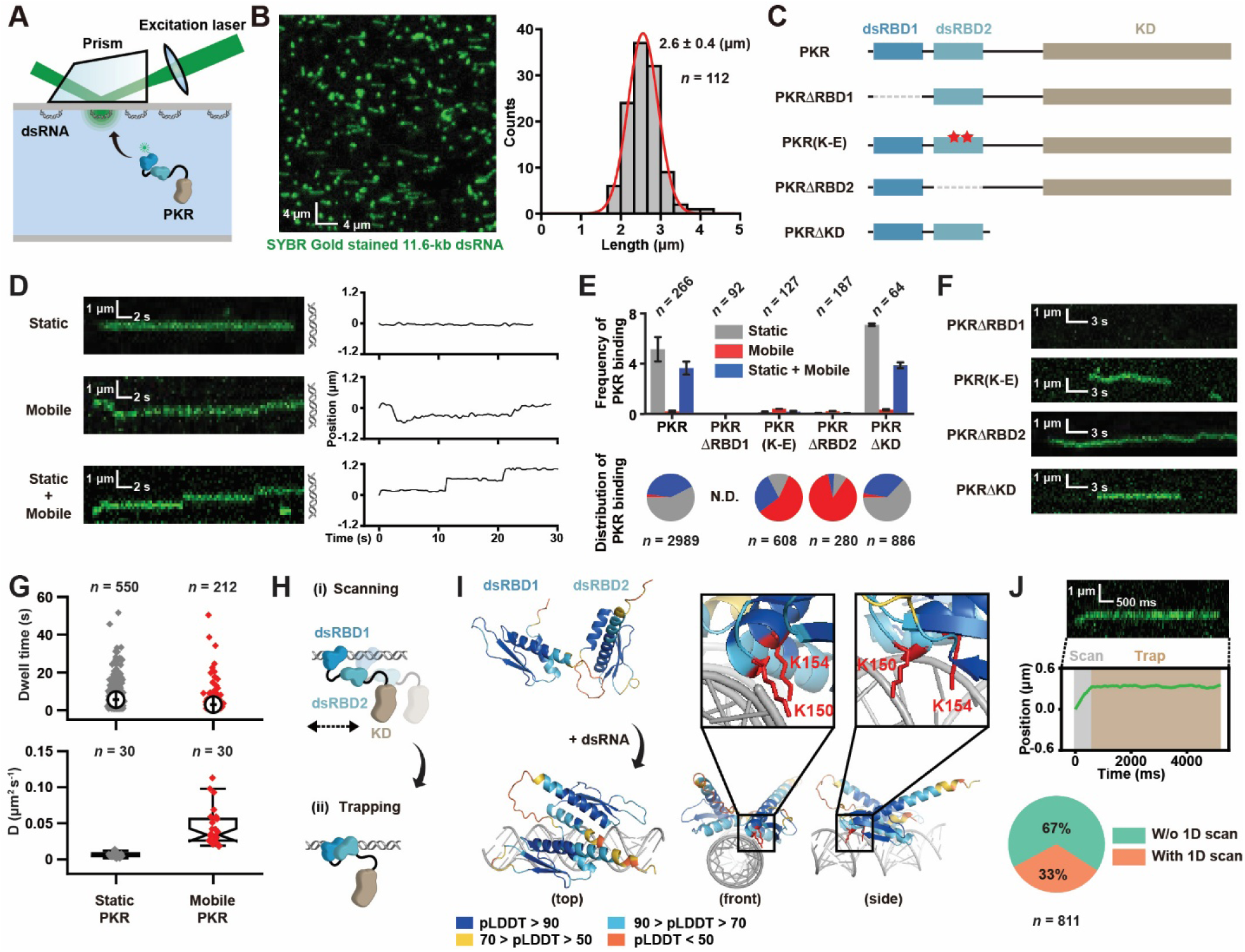
PKR switches between two RNA-binding modes. (**A**) A schematic illustration of dsRNA and PKR observations by prism-based smTIRF microscopy. (**B**) Left: Representative 11.6-kb dsRNA visualized by smTIRF microscopy in the absence of flow. The dsRNA was stained with SYBR Gold and a 42.7 x 42.7 µm field of view is shown. Right: the length distribution of the 11.6-kb dsRNA observed by smTIRF microscopy (*n* = number of dsRNA molecules). The data was fit with a Gaussian distribution that determined the mean ± **s.d.**. The contour length of an 11.6-kb dsRNA is calculated to be 3.3 µm. (**C**) A schematic illustration of PKR and PKR variants used in single-molecule studies. (**D**) Representative kymographs (left) and trajectories (right) showing two different binding modes of PKR on 11.6-kb dsRNA. (**E**) Top: Frequency of 20 pM PKR, PKRΔRBD1, PKR(K-E), PKRΔRBD2 and PKRΔKD binding to 11.6-kb dsRNA (mean ± s.d., *n* = number of dsRNA molecules). Bottom: Pie charts showing the distributions of binding events (*n* = total number of events; N.D.: not detected). (**F**) Representative kymographs showing the binding of 20 pM PKRΔRBD1, PKR(K-E), PKRΔRBD2, and PKRΔKD on 11.6-kb dsRNA. (**G**) Dwell time distributions (top) and box plots of diffusion coefficient (*D*, bottom) for static and mobile PKR on 11.6-kb dsRNA (*n* = number of events). Diamonds represent individual events. (**H**) A schematic illustration showing the “scan-and-trap” model of PKR binding to 11.6-kb dsRNA. (**I**) AlphaFold models of PKR’s dsRBDs in the absence (top) or presence (bottom) of a 25-bp dsRNA. The closeups show the key contacts between dsRBD2 and dsRNA. (**J**) Top: Representative kymograph and trajectory showing the “scan-and-trap” process at 50-ms imaging intervals. Bottom: Pie chart showing the distributions of static PKR binding with and without 1D scanning at 50-ms imaging intervals (n = number of binding events).

To identify the domains responsible for these binding modes, we separately deleted dsRBD1, dsRBD2, and the KD, generating three truncated proteins PKRΔRBD1, PKRΔRBD2, and PKRΔKD (**Figures 1C and S1B**). Additionally, an RNA-binding mutant in dsRBD2 (K150E and K154E), refer to as PKR(K-E), was introduced to abolish dsRBD2-RNA interactions but retain the potential regulatory roles of dsRBD2^27,36–38^ (**Figures 1C and S1B**). These variants were then assessed for their binding to dsRNA at the single-molecule level. Deletion of dsRBD1 removed all PKR binding to 11.6-kb dsRNA, consistent with its determinative role in RNA recognition^39,40^ (**Figures 1E and 1F**). In contrast, PKR(K-E) and PKRΔRBD2 exhibited highly similar profiles: although both retained the mobile binding, their static binding was nearly eliminated (**Figures 1E and 1F**). These findings indicated that: 1) both dsRBDs contribute to PKR-dsRNA binding, with dsRBD1 responsible for mobile binding and dsRBD2 for static binding; 2) mobile binding is required for static binding. Notably, deletion of the KD had no effect on PKR binding modes or frequency, suggesting that the KD is not involved in PKR-dsRNA interactions at the single-molecule level (**Figures 1E and 1F**).

The dwell times of static and mobile PKR on 11.6-kb dsRNA were 5 seconds (s) and 3 s, respectively (**Figure 1G**, top). The observed diffusion coefficient of static PKR was 0.006 μm^2^ s^-1^, likely reflecting the fluctuation of dsRNA under smTIRF. In contrast, the diffusion coefficient of mobile PKR was 0.0428 μm^2^ s^-1^, indicating actual movement (**Figure 1G**, bottom). Together, these findings appear to support a two-step capture model for PKR-RNA interaction (**Figure 1H**). Initially, PKR utilizes dsRBD1 to scan the dsRNA through passive 1D sliding. dsRBD2 then traps this diffusive complex on the RNA to establish static binding, a process we refer to as “scan-and-trap”. Consistent with this model, AlphaFold predictions suggested that the two dsRBDs come into a close proximity upon the dsRNA binding, with two lysine residues (K150, K154) in dsRBD2 helping position PKR on the dsRNA (**Figure 1I**). To validate this further, we reduced the imaging intervals to 50 millisecond (ms) and found that 33% of static binding events were preceded by a 1D scanning (**Figures 1J and S1D**). Since the dwell time for these 1D scanning events was near the 50-ms time resolution limit of our system (τ_on·1D scan_= 66 ms, **Figure S1E**), we envisioned that the “scan-and-trap” process might occur too quickly for the 1D scanning to be recorded in the remaining 67% of events.

### dsRBD2 compensates dsRBD1’s limitation on short dsRNAs

The role of dsRBD1 in recruiting dsRBD2 onto dsRNA was further illustrated by comparing the binding affinity of PKR and PKRΔRBD1 to 11.6-kb dsRNA across a range of protein concentrations (**Figures 2A and 2B**). The observed difference exceeded four orders of magnitude, supporting that dsRBD2 has no inherent RNA-binding activity, but depends on dsRBD1 for functionality (**Figures 2A and 2B**).

**Figure 2.**
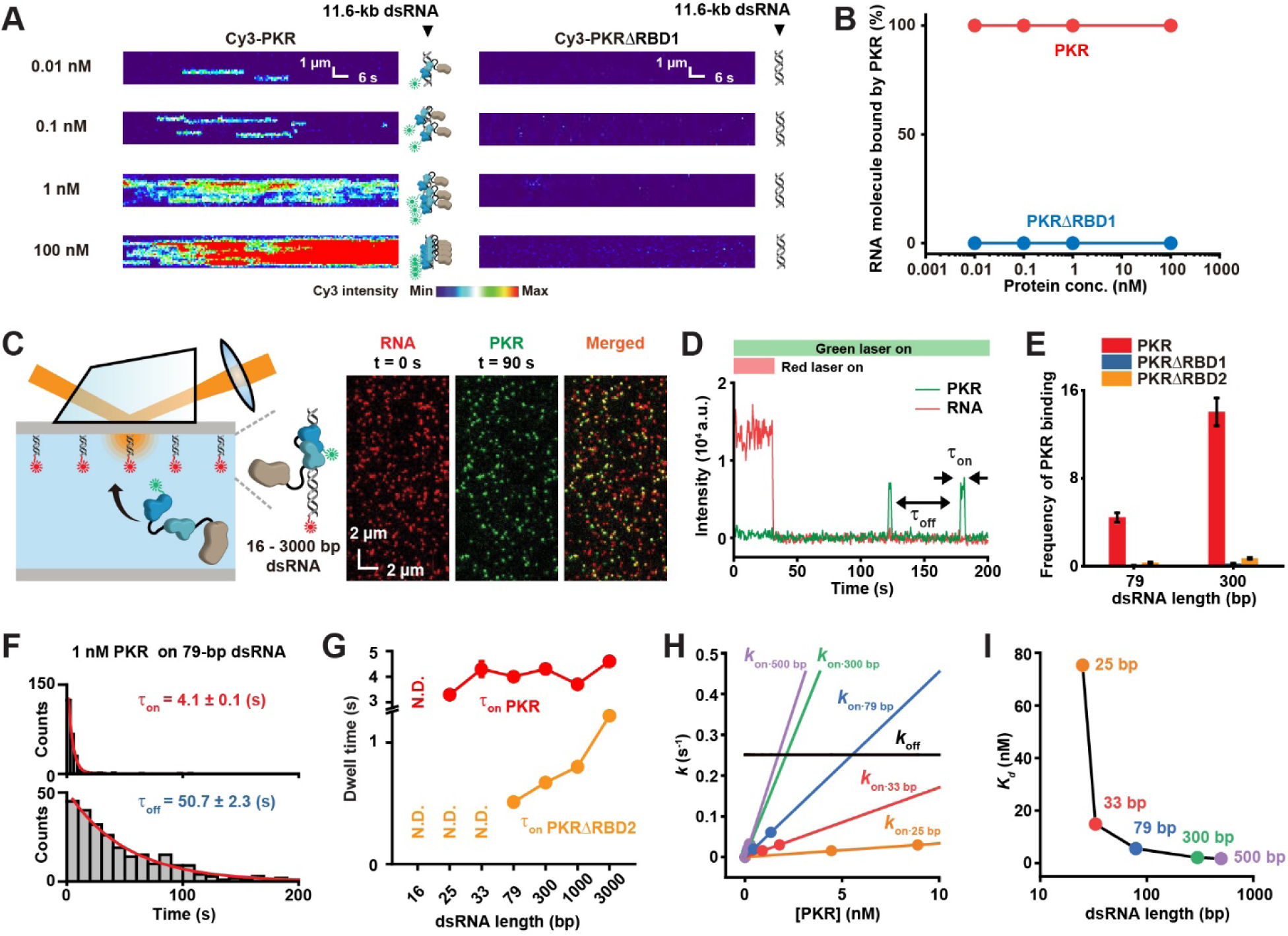
dsRBDs complement each other in shaping PKR binding kinetics. (**A**) Representative kymographs showing that Cy3-PKR binds to the 11.6-kb dsRNA, whereas Cy3-PKRΔRBD1 fails to bind, across a range of protein concentrations. dsRNA positions are indicated to the right of each kymograph. Fluorescent intensities were normalized to generate the heatmaps. (**B**) Statistics on the percentage of dsRNA molecules bound by PKR or PKRΔRBD1 at varying protein concentrations. (**C**) A schematic illustration (left) and representative images (right) of two-colored single-molecule co-localization assay. Cy5-dsRNAs are shown in red and Cy3-PKR is shown in green. Merged images were generated by overlaying both channels. (**D**) Representative trajectories showing two PKR binding events observed by single-molecule colocalization. τ_on_ and τ_off_ represent the on- and off-time of PKR binding, respectively. (**E**) The frequency of 0.5 nM PKR, PKRΔRBD1 or PKRΔRBD2 binding to 79-bp or 300-bp dsRNA (mean ± s.d.). (**F**) τ_on_ (top) and τ_off_ (bottom) distributions for 1 nM PKR binding to 79-bp dsRNA. Data were fit to a single exponential decay to derive the average dwell time (mean ± s.e.). (**G**) τ_on_ (mean ± s.e.) for PKR and PKRΔRBD2 binding to various dsRNAs. N.D.: not detected, likely due to insufficient time resolution of imaging. (**H**) PKR concentration dependence of *k_on_*. Data were fit to linear functions to derive the slopes and intercepts. (**I**) The dissociation equilibrium constants (*K_d_*) of PKR binding to various dsRNAs. *K_d_* was calculated from the intercept where *k_on_* = *k_off_* in (**H**).

Next, to investigate whether dsRBD1 also recruits dsRBD2 to other dsRNAs, we employed a recently developed two-color colocalization assay^22^ to monitor Cy3-PKR binding to Cy5-labeled dsRNA of various lengths from 16 to 3,000 bp (**Figures 2C**, **2D**, **S2A and S2B**). Similar to our previous findings, numerous single PKR binding events were observed on short dsRNA^22^ and the vast majority of them required both dsRBD1 and dsRBD2, exemplified by the 79-bp and 300-bp dsRNAs shown in **Figure 2E**. These single-molecule trajectories were then tracked to determine the on-times (τ_on_) (**Figures 2D**, **2F and S2B**). Of note, while stable binding event was rarely detected on 16-bp dsRNA, PKR displayed a consistent dwell time (τ_on PKR_ ≈ 4 s) on dsRNAs ranging from 25 to 3,000 bp (**Figure 2G**), nearly identical to that of static PKR on the 11.6-kb dsRNA (τ_on_ = 5 s, **Figure 1G**, top). In contrast, the observed dwell times of PKRΔRBD2 on these dsRNAs increased with the dsRNA length (τ_on PKRΔRBD2_ = 0.5–1.5 s, **Figures 2G and S2C**) but were much shorter than that of mobile PKR on 11.6-kb dsRNA (τ_on_ = 3 s, **Figure 1G**, top). These results supported the notion that sliding PKRΔRBD2 rapidly dissociated from short dsRNAs at the open end, whereas dsRBD2 trapped and stabilized PKR on these ligands.

The off-times (τ_off_) of single PKR binding events were measured across various protein concentrations (**Figures 2D**, **2F and S2D**), enabling the calculation of the association rate constant (*k_on_*) as 1/τ_off_ (**Figure 2H**). With the unchanged dissociation rate constant (*k_off_*, 1/τ_on_), we determined the dissociation constant (*K_d_*) for PKR binding (**Figures 2H**, **2I**). Notably, *k_on_* increased markedly with the ligand length (**Figure 2H**), while *K_d_* decreased substantially (**Figure 2I**). These findings are consistent with the idea that the 1D scanning mode is the rate-limiting step for static binding, as longer dsRNAs extend the 1D scanning lifetime (**Table S3**), enhancing the likelihood of being captured by dsRBD2. Together, these observations suggest that dsRBD1 has a high dissociation rate due to sliding, while dsRBD2 cannot bind independently; however, their coordinated action compensates for each other’s limitations, enabling high-affinity binding even on short dsRNAs.

### Prolonged lifetime triggers PKR dimerization and activation

While dsRBD2 extends PKR lifetimes on short dsRNAs from milliseconds to several seconds (**Figure 2G**), this static complex may function as an essential scaffold for KD dimerization. To examine this, we increased the PKR protein concentration in the two-color colocalization assay, allowing multiple PKR molecules to bind to a single dsRNA (**Figure 3A**). We found that PKR recognized the majority of dsRNA molecules under this condition, with only a small fraction of RNAs showing no binding events (no binding, N, **Figures 3A and 3B**). Three distinct types of PKR binding patterns were observed. The first type involved multiple single-molecule binding events, where each event was clearly separated in time (single binding, S, **Figure 3A**). The second type consisted of numerous independent PKR association–dissociation events that overlapped in time, leading to rapid fluctuations in Cy3 fluorescence intensity that quickly returned to background levels (multiple binding, M, **Figure 3A**). The third type showed several steps of PKR accumulation, followed by stable signals that lasted for hundreds of seconds, likely representing PKR dimerization/polymerization (polymerization, P, **Figure 3A**).

**Figure 3.**
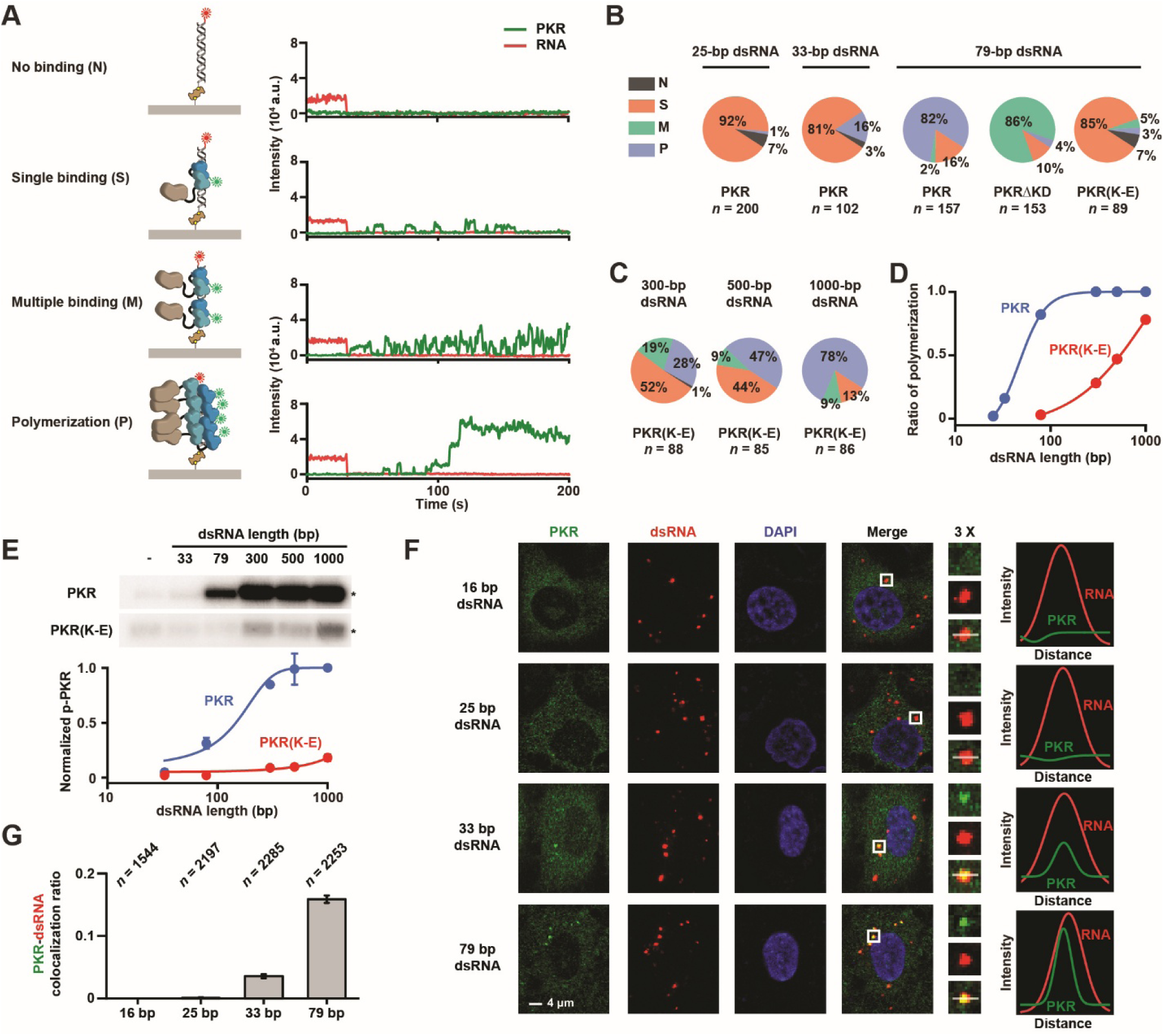
dsRBD2 promotes KD dimerization by extending binding lifetime. (**A**) Illustrations and representative trajectories showing three distinct binding patterns of PKR (10 nM) on dsRNA. (**B**) Pie charts showing the distributions of PKR, PKRΔKD and PKR(K-E) (10 nM) binding event on various dsRNAs (*n* = total number of dsRNA molecules examined). (**C**) Pie charts showing the distributions of PKR(K-E) (10 nM) binding event on dsRNAs of varying length (*n* = total number of dsRNA molecules examined). (**D**) The ratio of PKR (10 nM) and PKR(K-E) (10 nM) polymerization event on dsRNAs of varying length. (**E**) Top: *In vitro* kinase phosphorylation assay showing the activation of PKR and PKR(K-E) (200 nM) by various dsRNAs (2 mg/L). Stars indicate phosphorylated PKR (p-PKR). Bottom: Quantitative analysis of p-PKR (mean ± s.d.). (**F**) Representative IF images showing endogenous PKR colocalized with 33-bp and 79-bp dsRNA, but not with 16-bp and 25-bp dsRNA in A549 cells. Enlarged views of selected foci (indicated by boxes) are shown with corresponding relative fluorescence intensity profiles (line drawing in the enlarged images). (**G**) The ratio of immunofluorescence (IF) colocalization between endogenous PKR and various Cy3-dsRNAs in A549 cells (mean ± s.d.; *n* = total number of dsRNA foci examined).

Nearly all PKR bound to 25-bp dsRNA in singular, while dimerization was occasionally detected on 33-bp dsRNA (**Figure 3B**). Notably, a PKR dimer exhibited extremely long binding dwell times, often remaining bound until the end of single-molecule recording (**Figure S3A**). In contrast to the 25- and 33-bp dsRNAs, 82% of PKR binding on 79-bp dsRNA involved dimerization or polymerization, with 2% corresponding to multiple binding and 16% to single binding events (**Figure 3B**). We substituted the *wild-type* PKR with PKRΔKD and found that most dimerization/polymerization events on 79-bp dsRNA were turned into multiple binding events (**Figures 3B and S3B**), further confirming that these stable PKR binding events were KD-dependent. Replacing PKR with PKR(K-E) also caused a deficiency in KD dimerization/polymerization, but with most interactions resulting in singular binding (**Figures 3B and S3B**). These observations suggest that the dsRBD2-RNA interactions play a key role in promoting KD-KD interactions.

The lack of KD dimerization/polymerization for PKR(K-E) on 79-bp dsRNA is likely due to the short-lived dsRBD1-RNA interactions, where the first PKR(K-E) scanned along dsRNA and quickly dissociated before the next one could bind. To test this hypothesis, we measured PKR and PKR(K-E) binding events on dsRNAs of varying lengths, ranging from 25 bp to 1000 bp (**Figures 3C, 3D and S3C**). Remarkably, the ratio of dimerization/polymerization events increased with the dsRNA length, with PKR forming oligomers much more efficiently than PKR(K-E) (**Figures 3C and 3D**). Nevertheless, longer dsRNAs (> 300 bp) were able to trigger KD-KD interactions between PKR(K-E) molecules, consistent with the notion that dimerization/polymerization probability is increased by the extended dwell times (**Figures 3C and 3D**). These findings demonstrate that the absence of a functional dsRBD2 can be compensated by a longer binding dwell time and the principal role of dsRBD2 is to ensure a prolonged lifetime on short dsRNAs. The significance of this time-limiting step was further validated by an *in vitro* kinase phosphorylation assay using various dsRNAs, where PKR(K-E) also required longer duplex for activation (**Figure 3E**).

To further investigate the kinetics of PKR-dsRNA binding within cells, we stimulated A549 cells with Cy3-labeled dsRNA of varying lengths and examined their colocalization with endogenous PKR using immunofluorescence (IF) (**Figures 3F and 3G**). Stable PKR-dsRNA interactions are expected to produce colocalized foci^41^, whereas short-lived interactions are not. Following transfection with 16-bp or 25-bp dsRNA, ligands to which the kinase either did not stably bind or bound in singular (**Figures S2B and 3B**), PKR exhibited a uniform distribution in cytoplasm with no colocalization with these RNAs (**Figures 3F, 3G and S3D**). In contrast, transfections with 33-bp and 79-bp dsRNA led to the formation of PKR foci that always colocalized with these exogenous RNAs, with the frequency of colocalization increasing with the dsRNA length (**Figures 3F, 3G and S3D**). These observations align with our *in vitro* results, supporting the idea that interactions between a single PKR and dsRNA are short-lived, and that KD dimerization plays a key role in stabilizing these interactions.

### The dsRBD interplay enables PKR to proofread RNA

The findings above illustrate how the dsRBDs work together to recognize homoduplex RNAs that mimic viral RNA; however, whether this interplay extends to other RNAs remains unknown. To investigate this, we replaced the dsRNAs with a 1.8-knt Cy5-labeled 18S rRNA in the two-color colocalization assay (**Figures 4A and S4A**). This relatively long ssRNA can adopt complex secondary structures *in vitro*^42^, potentially allowing multiple PKR molecules to bind. Nevertheless, when dsRBD1-dependent PKR binding events were detected on the 18S rRNA, they were highly transient (half-life, t_1/2_ = 0.4 s) and did not lead to KD dimerization or polymerization (**Figures 4A and 4B**). Notably, PKR and PKRΔRBD2 exhibited equivalent binding frequencies and half-lives on the 18S rRNA (**Figures 4A and 4B**), suggesting that the dsRBD2 was not engaged with this ligand. As a control, a functional dsRBD2 significantly increased the half-life of single PKR binding to 79-bp dsRNA (**Figure 4B**), further demonstrating the absence of the dsRBD interplay on 18S rRNA.

**Figure 4.**
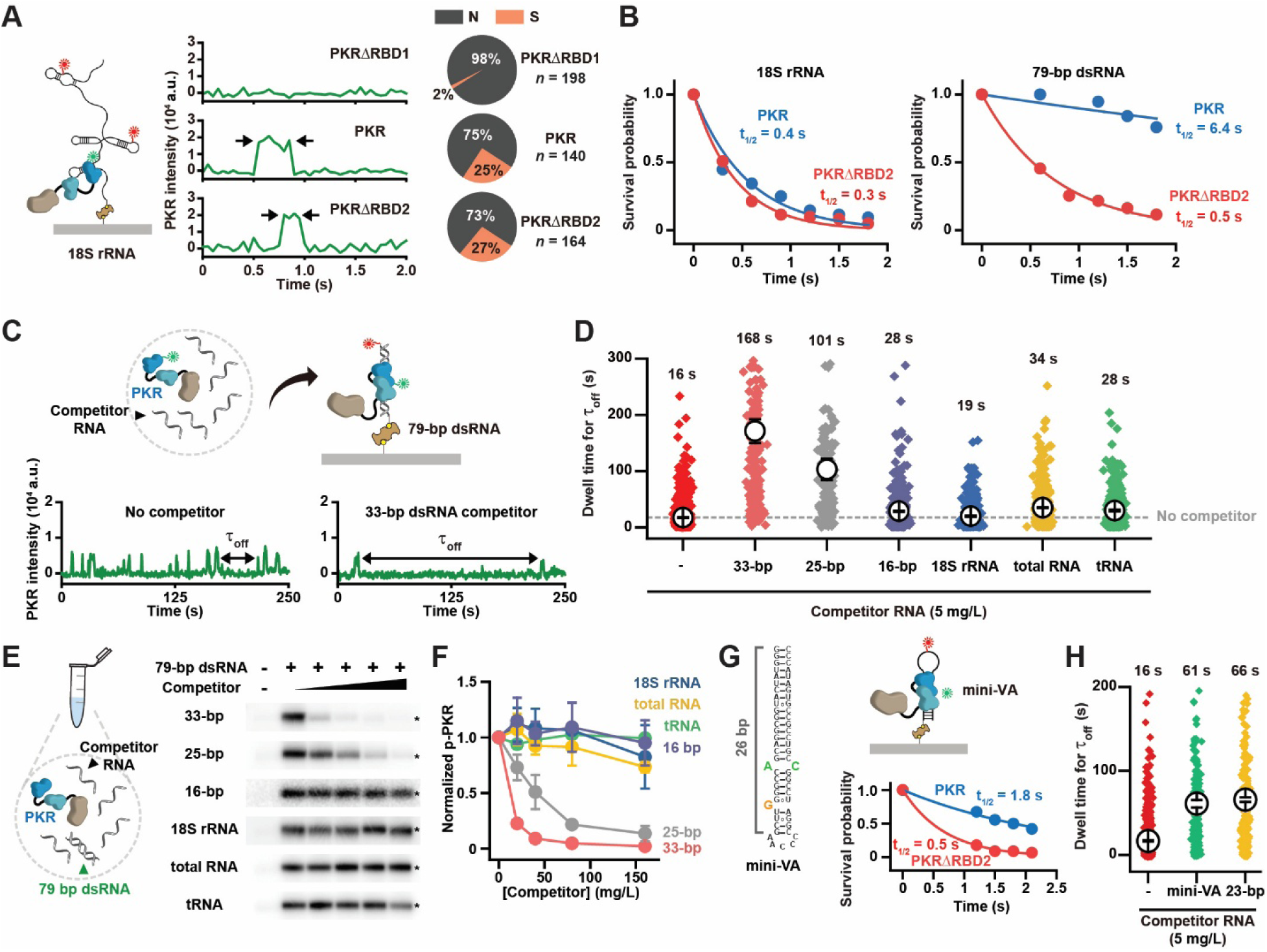
PKR proofreads RNA ligands through dsRBD interplay. (**A**) Left: A schematic illustration of Cy3-PKR binding to Cy5-labeled 18S rRNA measured by two-colored single-molecule co-localization assay. Center: Representative trajectories showing 10 nM Cy3-PKRΔRBD1, Cy3-PKR and Cy3-PKRΔRBD2 binding to 18S rRNA. Arrows indicate protein associations and dissociations. Right: Pie charts showing the distributions of 10 nM PKRΔRBD1, PKR and PKRΔRBD2 binding event on 18S rRNA (*n* = total number of RNA molecules examined). (**B**) Survival probability for PKR and PKRΔRBD2 on 18S rRNA and 79-bp dsRNA. Data were fit by exponential decay functions to obtain survival half-lives. (**C**) Top: A schematic illustration of the competition assay. PKR (3 nM) was incubated with 5 mg/L of various RNA competitors before injection to assess its binding to 79-bp dsRNA. Bottom: Representative trajectories showing the 33-bp dsRNA competitor extended the τ_off_ of PKR binding to 79-bp dsRNA. (**D**) τ_off_ distributions for 3 nM PKR binding to 79-bp dsRNA in the presence of various RNA competitors (5 mg/L). Diamonds represent individual events and open circles represent the mean dwell time derived by single exponential decay fitting (mean ± s.e.). (**E**) Left: A schematic illustration of *in vitro* kinase phosphorylation assay in the presence of RNA competitors. Right: PKR (200 nM) activation by 79-bp dsRNA (2 mg/L) with various RNA competitors (0, 20, 40, 80, 160 mg/L). Stars indicate phosphorylated PKR (p-PKR). (**F**) Quantitative analysis of p-PKR (mean ± s.d.). (**G**) Left: Secondary structure of mini-VA. Right: A schematic illustration of Cy3-PKR binding to Cy5-labeled mini-VA and survival probability for PKR and PKRΔRBD2 on mini-VA. Data were fit by exponential decay functions to obtain survival half-lives. (**H**) τ_off_ distributions for 3 nM PKR binding to 79-bp dsRNA in the presence of mini-VA and 23-bp dsRNA (5 mg/L). Diamonds represent individual events and open circles represent the mean dwell time derived by single exponential decay fitting (mean ± s.e.).

To validate this notion, we examined PKR binding to 79-bp dsRNA in the presence of excess amounts of various competitor RNAs (5 mg/L, **Figures 4C**, **4D and S4B**). Under conditions that allow the dsRBD interplay to occur, PKR became increasingly associated with these competitors, lowering the pool of free PKR and extending the off-times between binding events on the 79-bp dsRNA (**Figure 4C**). As expected, competitors like 33-bp and 25-bp dsRNAs that engage with dsRBD2, significantly increased the off-times by 10- and 6-fold, respectively (τ_off_ = 16 s, without competitor; τ_off_ = 168 s, with 33-bp dsRNA; τ_off_ = 101 s, with 25-bp dsRNA; **Figures 4D and S4C**). In contrast, 18S rRNA, total RNA, tRNA, and 16-bp dsRNA had a negligible impact on off-times (**Figures 4D and S4C**). In align with the single-molecule observations, 33-bp and 25- bp dsRNAs effectively blocked PKR phosphorylation induced by 79-bp dsRNA, whereas other competitors failed to inhibit PKR activation *in vitro*, even at substantially higher concentrations (**Figures 4E**, **4F**). These findings demonstrate that although PKR could bind RNAs like 18S rRNA and 16-bp dsRNA—as reflected by increasing off-times with rising competitor concentrations (**Figure S4D**)—their interactions were transient and limited to dsRBD1. The dsRBD interplay appeared to require a minimum dsRNA length of ∼22 bp, as τ_off_ remained unaffected below this threshold (**Figure S4E**).

We next explored PKR binding to the mini-VA sequence, a characterized PKR inhibitor described in the literature^43^ (**Figure 4G**). Although this 26-bp stem-loop structure is not a perfect duplex, the two dsRBDs work together in a manner similar to that observed on 79-bp dsRNA, with dsRBD2 significantly extending the half-life of PKR binding (**Figure 4G**). Consistent with its inhibitory role^43^, mini-VA increased the off-times by fourfold in the competition assay, though this effect was comparable to that of a 23-bp homoduplex (**Figure 4H**). Collectively, these observations suggest that the dsRBD interplay allows PKR to distinguish its ligands from the abundant non-ligand RNAs. This likely reflects the asymmetric binding kinetics between the two binding steps, which enables PKR to disregard numerous transient scanning events on non-ligand RNAs (such as rRNA and tRNA), while selectively responding to the static binding events on correct targets like 79-bp dsRNA and mini-VA.

### dsRBDs interrogate duplex integrity through their interplay

Although PKR can accommodate certain imperfect secondary structures (**Figure 4G**), bulges within an RNA duplex have been shown to hinder the kinase binding and activation^43–45^. This suggests that the dsRBDs must recognize essential characteristics of these imperfections, including their size, position, and type. To examine this, we introduced tandem 1- or 2-nt bulges or internal loops into the 79-bp dsRNA and assessed PKRΔRBD2 binding to these RNAs (**Figures 5A and 5B**). While imperfect structures decreased PKRΔRBD2 binding frequency in a size-dependent manner, no notable differences were found between dsRNAs with bulges and those with internal loops (**Figure 5C**). These results suggest that both structures have a mild impact on PKR scanning, likely by partially disrupting the interactions between dsRBD1 and dsRNA, without completely obstructing the 1D motion.

**Figure 5.**
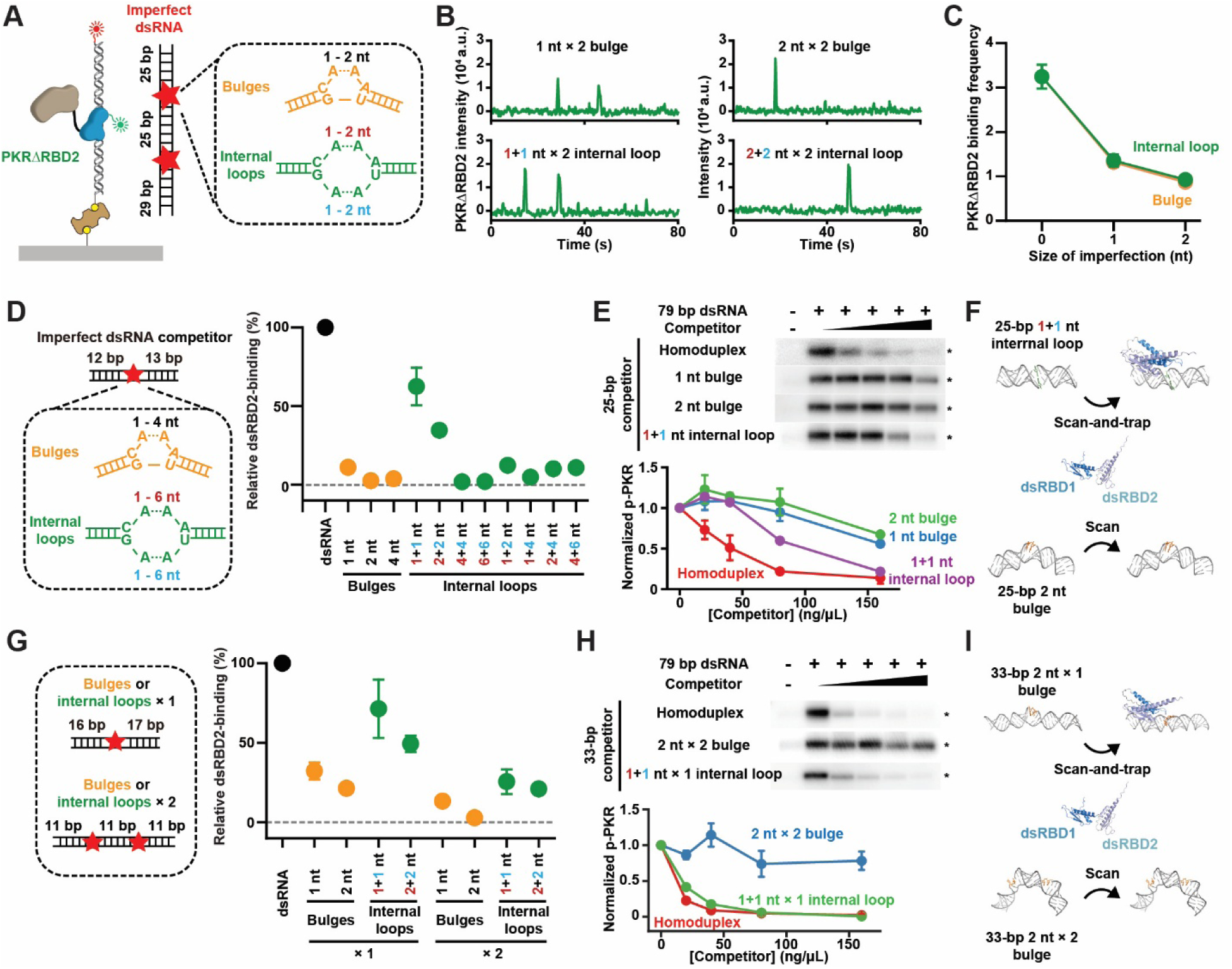
Bulges or large internal loops disrupt dsRBD interplay. (**A**) A schematic illustration of Cy3-PKRΔRBD2 binding to imperfect Cy5-labeled 79-bp dsRNA. (**B**) Representative trajectories showing PKRΔRBD2 binding to various imperfect 79-bp dsRNAs. (**C**) Frequency of PKRΔRBD2 binding to various imperfect 79-bp dsRNAs. (**D**) Left: A schematic illustration of imperfect 25-bp dsRNAs. Right: Relative binding affinity of PKR to imperfect 25-bp dsRNAs. (**E**) Top: PKR (200 nM) activation by 79-bp dsRNA (2 mg/L) with various imperfect 25-bp dsRNA competitors (0, 20, 40, 80, 160 mg/L). Stars indicate phosphorylated PKR (p-PKR). Bottom: Quantitative analysis of p-PKR (mean ± s.d.). (**F**) AlphaFold models showing that dsRBDs interacts with each other on a 25-bp dsRNA containing a small internal loop, but not on a 25-bp dsRNA containing a bulge. (**G**) Left: A schematic illustration of imperfect 33-bp dsRNAs. Right: Relative binding affinity of PKR to imperfect 33-bp dsRNAs. (**H**) Top: PKR (200 nM) activation by 79-bp dsRNA (2 mg/L) with various imperfect 33-bp dsRNA competitors (0, 20, 40, 80, 160 mg/L). Stars indicate phosphorylated PKR (p-PKR). Bottom: Quantitative analysis of p-PKR (mean ± s.d.). (**I**) AlphaFold models showing that dsRBDs interact with each other on a 33-bp dsRNA containing one bulge, but not on a 33-bp dsRNA containing two bulge in tandem.

We next introduced either a 1- to 4-nt bulge or a 1- to 6-nt internal loop at the center of a 25-bp dsRNA and evaluated dsRBD2 binding to these RNAs using the competition assay (**Figures 5D**, **S5A**). Binding was quantified relative to the τ_off_ observed with a 25-bp homoduplex competitor, which was set to 100% (**Figures 5D**, **S5B**). Remarkably, the presence of a single bulge of any size completely abolished dsRBD2 binding (**Figures 5D**, **S5B**). In contrast, small symmetric internal loops (1 or 2 nt on both strands) only modestly reduced binding affinity, whereas larger or asymmetric internal loops produced inhibitory effects comparable to a single bulge (**Figures 5D**, **S5B**). These differences in binding were consistent with the corresponding abilities of these RNAs to inhibit kinase activation *in vitro* (**Figure 5E**), reinforcing the idea that the “scan-and-trap” mechanism is sensitive to duplex integrity at the single base-pair resolution (**Figure 5F**). When the dsRNA length was increased to 33 bp, the inhibitory effect of a single structural feature remained evident, though less pronounced compared to the 25-bp dsRNA (**Figures 5G, 5H, S5C and S5D**). Notably, introducing tandem bulges or internal loops substantially amplified the inhibitory effect (**Figures 5G, 5H and 5I**). Collectively, these observations imply that dsRBD2 is selectively sequestrated from self-RNAs due to the prevalent presence of bulges and internal loops within their duplex regions.

## DISCUSSION

The dimerization of PKR’s KD has been considered a mechanism for distinguishing between self- and nonself-RNAs^11^. While our observations support this model, single-molecule analysis reveals a precedent step and an extra layer of discrimination indeed governed by the dsRBD interplay (**Figure 6**). Structural features inherent to self-RNAs create an energetically unfavorable landscape that discourages PKR occupation. When the RNA adopts viral-like structures, dsRBD2 captures the dsRBD1–RNA complex for 4 seconds, increasing the likelihood of occupancy and establishing a scaffold for KD–KD interactions. Dimerization or polymerization of KD then generates multivalent contacts with the dsRNAs, further prolonging residence time and increasing occupation probability (**Figure 6**). This cascading framework amplifies RNA discrimination by several orders of magnitude, equipping PKR with the precision needed to pinpoint rare viral RNAs within an overwhelming excess of self-RNAs.

**Figure 6.**
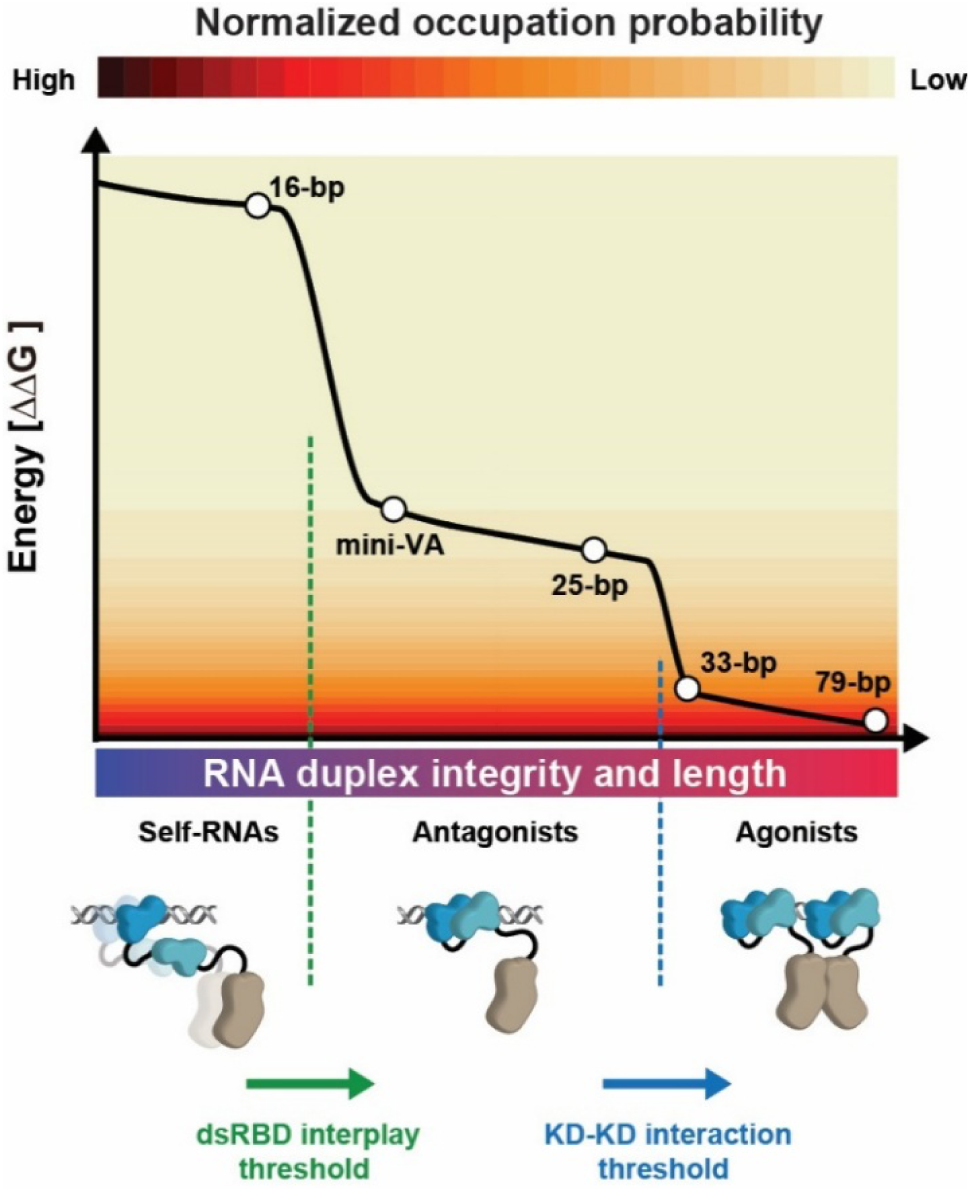
Energy landscape and a proofreading model for PKR sampling RNA ligands. PKR evaluates the integrity and length of the RNA duplex through the dsRBD interplay and the subsequent KD dimerization. As a monomer, PKR samples RNAs via dsRBD1, resulting in a low occupation probability on self-RNAs. When the dsRNA is sufficiently long and contains fewer imperfections, dsRBD2 stabilizes the dsRBD1–RNA complex in a static state lasting ∼4 s, which increases occupation probability and readily facilitates intermolecular KD–KD interactions. In the presence of two or more PKR molecules, dimers or polymers form and establish multivalent contacts with the RNA, substantially extending residence time and occupation probability. Of note, the full activation may depend on dimer–dimer interactions, supported by the notion that 33-bp dsRNA—despite strong binding—can act either as a weak agonist or as an antagonist of PKR. The heat map and open circles represent values for normalized occupation probability and ΔΔG values derived from experimental measurements and estimates. The black line is a representation of the landscape.

Multiple RBDs often cooperate, where binding of one domain enhances the effective concentration of nearby RBDs^1,8^. Tandem dsRBDs involved in endogenous dsRNA processing, such as those in TRBP, R3D1-L, and Staufen, have been shown to drive protein 1D diffusion by cooperative binding^5,46^. However, our data suggest that PKR functions through a distinct mode involving direct interactions between its dsRBDs. This aligns with the observations that PKR’s two dsRBDs originated from an evolutionary duplication event but have diverged in function^39,47^, and that dsRBD2 lacks the conserved GPxH (where x is any amino acid) RNA-binding motif^48–50^ (**Figure S5E**). Similar non-canonical dsRBDs are also found in other dsRBPs^48–50^, which may potentially contribute to controlling their binding kinetics. Collectively, these results reconcile the regulatory role of dsRBD2 in PKR activation–a function attributed to modulating kinase activity in earlier studies^27,36,51,52^.

Our observations demonstrate that PKR stoichiometry on RNA is determined not only by footprint but also by the binding mode. The detection of a PKR dimer on 33-bp dsRNA implies that a single PKR could, in theory, attach to a 16-bp dsRNA through its dsRBD2. However, no interaction was observed between dsRBD2 and the 16-bp dsRNA (**Figure 2G**), consistent with a recent report^43^. This cannot be attributed to footprint constraints, as the combined footprint of two dsRBDs spans only ∼15 bp (**Figure 1I**)^25^. These findings suggest that “scan-and-trap” binding mode requires a longer duplex (∼ 25 bp) to allow efficient dsRBD1 sliding, which in turn facilitates dsRBD2 loading onto the RNA (**Figures 1H and S4E**). On 33-bp dsRNA, the first PKR occupies ∼15 bp along a half-axis, potentially leaving the other half-axis available for a second PKR to bind via the same mode (**Figure 1I**). This footprint also explains why dsRBD2 fails to stably associate with a 25-bp dsRNA containing a central bulge or a 33-bp dsRNA with two tandem bulges (**Figure 5**), and is consistent with dsRBD2’s ability to bind the mini-VA sequence, which carries a bulge near one end (**Figure 4G**).

Due to its versatile roles beyond antiviral defense, considerable efforts have been devoted to identifying PKR antagonists^53^, as its aberrant activity has been linked to various autoimmune and autoinflammatory conditions^14–20^. In particular, RNA decoys that imitate dsRNA have the potential to compete with the endogenous agonist and sequester PKR^21,22^. Our findings underscore a central challenge in developing PKR antagonists: the fleeting interactions between individual PKR molecules and their RNA ligands. The formation of a dimer can substantially stabilize these contacts, but it also carries the risk of triggering PKR activation. Therefore, an effective RNA antagonist may require efficient dsRBD interplay combined with restrained KD–KD interactions^22^, though conditions required to achieve this balance warrant further characterization. Furthermore, disruption of the proposed dsRBD interactions results in a substantial decrease in PKR autophosphorylation (**Figure 3**), also pointing to a previously unknown therapeutic strategy to target aberrant activation using small molecules or proteins.

## METHODS

### Plasmid construction, recombinant protein labeling and purification

Human PKR, PKRΔRBD1 (residues Δ8-84), PKRΔRBD2 (residues Δ100-176), PKR(K-E), and PKRΔKD (residues Δ185-551) were purified and labeled using sortase-mediated peptide ligation^35^. In brief, sequences encoding PKR and λ-PPase were cloned into a pET28a plasmid. A hexa- histidine (his_6_) tag and a sortase recognition sequence (srt, LPETG) were introduced at the N-terminus of PKR. The PKRΔRBD1, PKRΔRBD2, PKR(K-E), and PKRΔKD mutations were generated using the KOD-Plus-Mutagenesis Kit (Toyobo, cat. no. SMK-101). Two serine residues separated the his_6_ and srt sequences, and these tags were linked to the proteins via a GGS linker. All plasmid constructs were amplified in *E. coli* DH5α and verified by DNA sequencing.

After transformation with the PKR expression plasmid, a single colony of Rosetta 2(DE3) pLysS was diluted into 1 L of LB containing 50 μg/mL kanamycin. At OD_600_ = 0.5, the growth temperature was decreased to 25 °C, and protein expression was induced by the addition of IPTG (0.2 mM) at 25 °C for 16 h. Cells were harvested and resuspended in Freezing Buffer (25 mM HEPES pH 8.0, 500 mM NaCl, 10% glycerol and 20 mM imidazole). The cell pellets were freeze-thawed three times, sonicated twice, and centrifuged at 48,000 × g for 1 hour (h). The supernatant was then loaded onto a Ni-NTA column, washed with Buffer A (25 mM HEPES pH 8.0, 500 mM NaCl, 10% glycerol and 20 mM imidazole) and eluted with Buffer B (25 mM HEPES pH 8.0, 500 mM NaCl, 10% glycerol and 300 mM imidazole). Fractions containing proteins were pooled and dialyzed overnight against Buffer C (25 mM HEPES pH 8.0, 200 mM NaCl, 1 mM DTT and 10% glycerol). The dialyzed proteins were loaded onto a heparin column, washed with Buffer D (25 mM HEPES pH 8.0, 1 mM DTT, 10% glycerol and 0.1 mM EDTA) plus 200 mM NaCl and eluted with Buffer D plus 1 M NaCl. Fractions containing proteins were further purified by size-exclusion chromatography (SEC, Superdex 200 10/300 GL, GE Healthcare). The protein fractions were pooled and dialyzed overnight against Labeling Buffer (50 mM Tris-HCl pH 8.0, 500 mM NaCl, 10 mM CaCl_2_ and 10% glycerol). For labeling, PKR was incubated with sortase and Cy3-labeled peptides (Cy3-CLPETGG, purchased from ChinaPeptides Co.,LTD) at 4 °C for 1 h (protein: sortase: peptide in the ratio of 1:1:5). After labeling, the reaction mixture was loaded onto a Ni-NTA column. The flow-through fractions were collected and diluted with Buffer D to reduce NaCl concentration within the buffer (∼ 100 mM). These fractions were then loaded onto a heparin column, extensively washed with Buffer D plus 100 mM NaCl and eluted with Buffer D plus 1 M NaCl. Protein-containing fractions were dialyzed against Storage Buffer (25 mM HEPES pH 8.0, 500 mM NaCl, 1 mM DTT, 0.1 mM EDTA, and 10% glycerol) and frozen at - 80 °C. The PKR variants were purified with similar procedures.

The concentrations and labeling efficiencies of all recombinant proteins were determined by measuring protein absorbance at 280 nm and Cy3 absorbance at 550 nm (**Table S2**), respectively.

### Preparation of RNA ligands

Two complementary 11,600-nt ssRNA strands were produced by SP6 *in vitro* transcription, and annealed together with two biotin-labeled ssRNA linkers to generate an 11.6-kb dsRNA as described^54^. In brief, two plasmids containing the SP6 promoter and sense/anti-sense strand sequences were constructed, amplified in *E. coli* DH5α and verified by DNA sequencing. DNA templates were then linearized by KpnI (sense strand) or BamHI (anti-sense strand) digestion, followed by purification through phenol-chloroform extraction and ethanol precipitation. RNAs were transcribed *in vitro* using SP6 RNA Polymerase (Thermo Fisher Scientific) and further purified by phenol-chloroform extraction and ethanol precipitation. Sense and anti-sense RNA strands (100 nM for each), along with RNA linker 1 and RNA linker 2 (10 µM for each) were mixed in Annealing Buffer A (20 mM Tris-HCl pH 7.5, 50 mM NaCl, 1 mM EDTA), denatured at 95 °C for 10 min, then cooled down slowly to 23 °C. The resulting product was separated on a 0.75% agarose gel, and the 11.6-kb dsRNA was excised and purified using Agarose Gel DNA Extraction Kit (Takara).

To generate RNAs used in the two-color single-molecule co-localization assay, short homoduplexes like 16-bp, 25-bp, 33-bp, and 79-bp dsRNA were prepared by mixing complementary oligonucleotides (5 µM for each**, Table S1**) in Annealing Buffer A, denaturing at 60 °C for 10 min, then cooling down slowly to 23 °C. To generate the 300-bp, 500-bp, 1,000-bp, and 3,000-bp dsRNAs, two single-stranded RNAs (ssRNAs) were synthesized by *in vitro* transcription, then annealed with a 50-nt biotin- and Cy5-labeled RNA linker 3 (5 µM**, Table S1**). To generate the 18S rRNA used in two-colored single-molecule co-localization assay, plasmids containing the T7 promoter and 18S rRNA sequences were constructed. The DNA template was linearized by XhoI digestion, and purified by phenol-chloroform extraction and ethanol precipitation. Fluorescently labeled 18S rRNA was synthesized by incorporating Cy5-UTP (Enzo Lifesciences) during transcription. Transcribed RNAs were resolved on a denaturing urea polyacrylamide gel, stained with ethidium bromide, and excised for purification. Cy5-labeled 18S rRNA (50 nM) was then annealed with a biotin-labeled DNA linker 1 (5 µM, **Table S1**) in Annealing Buffer B (10 mM Tris-HCl pH 7.5, 5 mM MgCl_2_), denatured at 60 °C for 10 min, then cooled down slowly to 23 °C. Biotin and Cy5-labeled mini-VA (50 nM) was purchased from GenScript (China), denatured in Annealing Buffer B at 60 °C for 10 min, then cooled down slowly to 23 °C. Total RNA was extracted using RNAiso Plus reagent (Takara). tRNA was kindly provided by Xiao-Long Zhou’s lab.

### Single-molecule imaging buffers and experiment conditions

The single-molecule Imaging Buffer consisted of 20 mM Tris-HCl (pH 7.5), 0.1 mM DTT, 0.2 mg/mL acetylated BSA (Molecular Cloning Laboratories), 0.0025% P-20 surfactant (GE healthcare), 5 mM MgCl_2_ and 100 mM NaCl. All single-molecule experiments were conducted at 23 °C. To minimize photoblinking and photobleaching, all Imaging Buffer was supplemented with a photostability enhancing and oxygen scavenging cocktail containing saturated (∼3 mM) Trolox and PCA/PCD oxygen scavenger system composed of PCA (1 mM) and PCD (10 nM)^55^.

### Single-molecule total internal reflection fluorescence (smTIRF) microscopy

All the single-molecule total internal reflection fluorescence (smTIRF) data were acquired on a custom-built prism-type TIRF microscope established on the Olympus IX73 microscope body^56^. Fluorophores were excited using the 532 nm for green and 637 nm for red laser lines built into the smTIRF system. Image acquisition was performed using an EMCCD camera (iXon Ultra 897, Andor) after splitting emissions by an optical setup (OptoSplit II emission image splitter, Cairn Research). Micro-Manager image capture software was used to control the opening and closing of a shutter, which in turn controlled the laser excitation.

To examine PKR binding on 11.6-kb dsRNA, RNA ligand (50 pM) in 500 μL T50 buffer (20 mM Tris- HCl pH 7.5, 50 mM NaCl) was injected into a custom-made flow cell chamber and stretched by laminar flow (300 μL/min). The stretched dsRNA was anchored at both ends to a neutravidin-coated, PEG-passivated quartz slide surface, and the unbound dsRNA was flushed by similar laminar flow. The *wild-type* Cy3-PKR or PKR variants (20 pM) in Imaging Buffer were introduced into the flow cell chamber. Unless stated otherwise, a 300-ms frame rate was used for single-molecule imaging. The PKR molecules on dsRNA were then monitored in real-time for 7.5 min in the absence of flow. After real-time recording, the dsRNA was located by staining with SYBR Gold (0.2 X, Invitrogen).

To examine PKR binding to dsRNAs (16-bp, 25-bp, 33-bp, 79-bp, 300-bp, 500-bp, 1,000- bp and 3,000-bp) in two-colored single-molecule co-localization assay, Cy5-labeled dsRNA (15 pM) in 300 μL T50 buffer was injected into a custom-made flow cell chamber by laminar flow (25 μL/min), and the unbound dsRNA was flushed by similar laminar flow. The *wild-type* PKR or PKR variants (0.5 nM) in Imaging Buffer were introduced into the flow cell chamber. A 100-ms or 300- ms frame rate was used for single-molecule imaging. The PKR molecules on dsRNA were then monitored in real-time for 7.5 min. To examine the binding patterns of PKR or PKR(K-E) on dsRNAs (25-bp, 33-bp, 79-bp, 300-bp, 500-bp and 1,000-bp), 10 nM Cy3-PKR or Cy3-PKR(K-E) was introduced into the flow cell chamber by laminar flow (100 μL/min). The PKR or PKR(K-E) molecules on dsRNA were then monitored in real-time for 7.5 min.

To examine PKR or PKRΔRBD2 binding to 18S rRNA, a 50-ms frame rate was used to detect transient protein-RNA interactions. Cy5-labeled 18S rRNA (160 pM) in 300 μL T50 buffer was injected into the flow cell chamber by laminar flow (25 μL/min), and the unbound 18S rRNA was flushed by similar laminar flow. 10 nM Cy3-PKR or Cy3-PKRΔRBD2 in Imaging Buffer was introduced into the flow cell chamber by laminar flow (100 μL/min). The PKR or PKRΔRBD2 molecules on 18S rRNA were then monitored in real-time for 7.5 min.

To examine PKR binding to competitor RNAs, Cy5-labeled 79-bp dsRNA (15 pM) in 300 μL T50 buffer was injected into a custom-made flow cell chamber by laminar flow (25 μL/min), and the unbound dsRNA was flushed by similar laminar flow. 3 nM Cy3-PKR and 5 mg/L competitors were pre-incubated in Imaging Buffer for 3 min and then introduced into the flow cell chamber. The PKR molecules on dsRNA were then monitored in real-time for 7.5 min.

To examine PKR or PKRΔRBD2 binding to imperfect dsRNAs, a 100-ms frame rate was used. Cy5-labeled imperfect dsRNA (15 pM) in 300 μL T50 buffer was injected into the flow cell chamber by laminar flow (25 μL/min), and the unbound dsRNA was flushed by similar laminar flow. 10 nM Cy3-PKR or Cy3-PKRΔRBD2 in Imaging Buffer were introduced into the flow cell chamber. The PKR or PKRΔRBD2 molecules on dsRNA were then monitored in real-time for 5 min.

### *In vitro* kinase phosphorylation assay

To measure the phosphorylation of PKR and PKR(K-E) on dsRNA of different lengths, 200 nM proteins were incubated with 2 mg/L dsRNA in Activation Buffer (20mM HEPES pH 7.5, 100 mM NaCl, 4 mM MgCl_2_, 1 mM ATP, and 25 μCi [γ-^32^P] ATP) for 20 min at 30 °C. To determine the inhibition effects of competitors on PKR phosphorylation, 200 nM PKR was incubated with 2 mg/L 79-bp dsRNA and competitors in Activation Buffer for 20 min at 30 °C. Reactions were quenched by adding 2 × SDS Loading Buffer (50 mM Tris-HCl, 2% SDS, 0.1% Bromophenol blue, 10% glycerol, 100 mM DTT) and analyzed on 4-20% Bis-Tris PAGE. Gels were exposed to a storage phosphor screen, and band intensities of PKR or its variants were captured with an Amersham Typhoon RGB and quantified using ImageQuant.

### Data analysis of TIRF imaging

All kymographs were generated along the dsRNA by a kymograph plugin in ImageJ (J. Rietdorf and A. Seitz, EMBL Heidelberg). Particles were tracked using DiaTrack 3.05 to obtain single-molecule fluorescent intensities and trajectories. For studies involving Cy3-PKR and Cy5-labeled RNA ligands, fluorescent molecules in two channels were co-localized using a custom-written MATLAB script. PKR and RNA molecules were tracked by SPARTAN to generate trajectories^57^. For all analyses, signals with a lifetime shorter than three imaging frames were excluded.

To determine the frequency of PKR binding on dsRNA, single-molecule movies were recorded for 7.5 min. Proteins binding with a minimum lifetime of 0.3 s (for 100-ms frame rate) or 0.9 s (for 300-ms frame rate) were counted as the number of PKR binding (*N*_PKR-binding_). Following the real-time single-molecule recording, the number of dsRNA molecules (*N*_RNA_) was determined either by SYBR Gold staining or by Cy5 imaging. The frequencies of PKR binding (*F*_PKR-binding_) were calculated using the following equations, which also included corrections for labeling efficiencies of the proteins (the numbers in the denominator):

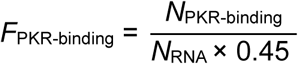

To determine the diffusion coefficients of PKR and PKRΔRBD2, particles were tracked using DiaTrack 3.05 to obtain single-molecule trajectories. Diffusion coefficients were calculated from the trajectories as previously described^35^. Briefly, the diffusion coefficient (*D*) was determined from the slope of a mean-square displacement (MSD) versus time plot using the equation MSD(t) = 2*D*t, where t is the time interval. The first 10% of the total measurement time was used for point fitting. A minimum number of 50 frames were used to calculate the diffusion coefficients.

To determine the binding pattern of PKR binding on dsRNA with various length, single-molecule movies were recorded for 7.5 min. Fluorescence intensity trajectories were extracted using SPARTAN. PKR binding events that were clearly separated in time were classified as single binding (S). Events where multiple PKR molecules bound in overlapping fashion but returned to background fluorescence within 50 seconds were categorized as multiple binding (M). Since a single PKR binding event typically lasted less than 50 seconds, events with stable fluorescence intensity and lifetimes exceeding 50 seconds were classified as polymerization (P). Events exhibiting both single binding and multiple binding during their course were categorized as multiple binding. Events exhibiting both multiple binding and polymerization during their course were categorized as polymerization.

### Immunofluorescence

A549 cells were seeded on cell confocal dishes (Cellvis) to reach 60 % confluency and transfected with Cy3-labeled dsRNA of varying lengths (16-bp, 25-bp, 33-bp and 79-bp; 500 mg/L) using Lipofectamine 2000 Reagent (Thermo Fisher Scientific). After fixation, A549 cells were incubated with rabbit anti-PKR IgG (Abways) followed by incubation with AlexaFluor488-conjugated donkey anti-rabbit IgG (Invitrogen) and DAPI. The cells were imaged using a Leica SP8 WLL confocal microscope in DAPI, AlexaFluor488, and Cy3 channels. Quantification of PKR and dsRNA colocalization was performed using ImageJ software.

## ACKNOWLEDGEMENT

This work was supported by the National Key R&D Program of China (2021YFA1300503), the Strategic Priority Research Program of the Chinese Academy of Science (XDB0570000) and the National Natural Science Foundation of China (32470928) to J. Liu.

## AUTHOR CONTRIBUTIONS

L.-L.C., and J.L. conceived the project; C.H., C.L., and J.L. designed the experiments; C.H., C.L., and Y.-F.X. purified the proteins and RNAs; C.H., and C.L. performed the single molecule studies; C.H., C.L. and M.R. performed the IF studies; C.H., C.L., Y.-F.X., and J.L. analyzed the data; C.H., C.L., L.-L.C., and J.L. wrote the paper and all authors participated in critical discussions.

## CONFLICT OF INTEREST

Authors declare no competing interests.

## Notes

### Competing Interest Statement

The authors have declared no competing interest.

