## Supplemental Figures and Tables for "Stepwise Ligand Capture Primes PKR for Activation and RNA Discrimination"

### Table S1. Oligonucleotides Used in This Study

| Name | Sequences |
|---|---|
| RNA linker 1 | GUACCGCGCUGAUAAGCCUGGUUGACGGAAGUGGCAAUUCUAGAGGGCCCUAUUC-biotin |
| RNA linker 2 | CCGGAUCGCUCGAGACGCAUUUGCAUCUAGAGGGCCCUAUUC-biotin |
| RNA linker 3 | Cy5-UGAGGAUCCCGGAUCGCUCGAGACGCAUUUGCAUCUAGAGGGCCCUAUUC-biotin |
| DNA linker 1 | Biotin-TCGAGTGGTCCAGGAACATTATCTGTCCCCTGTCAAATAGGGCCGCCTTATACC |
| 16-nt sense | Biotin-GGGUUUUCCCAGUCAC |
| 16-nt antisense | Cy5-GUGACUGGGAAAACCC |
| 25-nt sense | Biotin-GGGUUUUCCCAGUCACGACGUUGUA |
| 25-nt antisense | Cy5-UACAACGUCGUGACUGGGAAAACCC |
| 33-nt sense | Biotin-GGGUUUUCCCAGUCACGACGUUGUAAAACGACG |
| 33-nt antisense | Cy5-CGUCGUUUUACAACGUCGUGACUGGGAAAACCC |
| 79-nt sense | Biotin-GGGUUUUCCCAGUCACGACGUUGUAAAACGACGGCCAGUGAAUUCGAG-CUCGGUACCCGGGGAUCCUCUAGAGUCGACC |
| 79-nt antisense | Cy5-GGUCGACUCUAGAGGAUCCCCGGGUACCGAGCUCGAAUUCACUGGC-CGUCGUUUUACAACGUCGUGACUGGGAAAACCC |
| Mini-VA | Biotin-GGGUAUCAUGGCGGACGACCGGGGUUCGAACCCCGGAUCCGGCCGUCC-GCCGUGAUACCC-Cy5 |
| 25-nt RNA 1 | UACAACGUCGUGACUGGGAAAACCC |
| 25-nt-A RNA 1 | UACAACGUCGUGAACUGGGAAAACCC |
| 25-nt-2A RNA 1 | UACAACGUCGUGAAACUGGGAAAACCC |
| 25-nt-4A RNA 1 | UACAACGUCGUGAAAAACUGGGAAAACCC |
| 25 nt-6A RNA 1 | UACAACGUCGUGAAAAAAACUGGGAAAACCC |
| 25-nt RNA 2 | GGGUUUUCCCAGUCACGACGUUGUA |
| 25-nt-A RNA 2 | GGGUUUUCCCAGAUCACGACGUUGUA |
| 25-nt-2A RNA 2 | GGGUUUUCCCAGAAUCACGACGUUGUA |
| 25-nt-4A RNA 2 | GGGUUUUCCCAGAAAAUCACGACGUUGUA |
| 25-nt-6A RNA 2 | GGGUUUUCCCAGAAAAAAUCACGACGUUGUA |
| 33-nt RNA 1 | CGUCGUUUUACAACGUCGUGACUGGGAAAACCC |
| 33-nt-A RNA 1 | CGUCGUUUUACAACGUCAGUGACUGGGAAAACCC |
| 33-nt-2A RNA 1 | CGUCGUUUUACAACGUCAAGUGACUGGGAAAACCC |
| 33-nt-A×2 RNA 1 | CGUCGUUUUACAAACGUCGUGACAUGGGAAAACCC |

| | |
|---|---|
| 33-nt-2A×2 RNA 1 | CGUCGUUUUACAAAACGUCGUGACAAUGGGAAAACCC |
| 33-nt RNA 2 | GGGUUUUCCCAGUCACGACGUUGUAAAACGACG |
| 33-nt-A RNA 2 | GGGUUUUCCCAGUCACAGACGUUGUAAAACGACG |
| 33-nt-2A RNA 2 | GGGUUUUCCCAGUCACAAGACGUUGUAAAACGACG |
| 33-nt-A×2 RNA 2 | GGGUUUUCCCAAGUCACGACGUUAGUAAAACGACG |
| 33-nt-2A×2 RNA 2 | GGGUUUUCCCAAAGUCACGACGUUAAGUAAAACGACG |

**Table S2. Labeling Efficiencies of PKR and PKR variants**

| Protein | Labeling Efficiencies |
|---|---|
| Cy3 labeled PKR | 45% |
| Cy3 labeled PKRΔRBD1 | 44% |
| Cy3 labeled PKR(K-E) | 60% |
| Cy3 labeled PKRΔRBD2 | 70% |
| Cy3 labeled PKRΔKD | 33% |

**Table S3. Calculated $\tau_{on}$ and $k_{off}$ of PKR scanning on various dsRNAs**

| dsRNA Length (bp) | $\tau_{on}$ (ms)* | $k_{off}$ ($s^{-1}$) |
|---|---|---|
| 33 | 0.86 | 1162.78 |
| 79 | 4.93 | 202.90 |
| 100 | 7.90 | 126.63 |
| 300 | 71.07 | 14.07 |
| 500 | 197.43 | 5.07 |
| 1000 | 789.72 | 1.27 |

* The $\tau_{on}$ was calculated using the equation: $\tau_{on} = d^2/2D$, where d is the contour length of dsRNA and D is the average diffusion coefficient of mobile PKR (**Figure 1G**)[56].

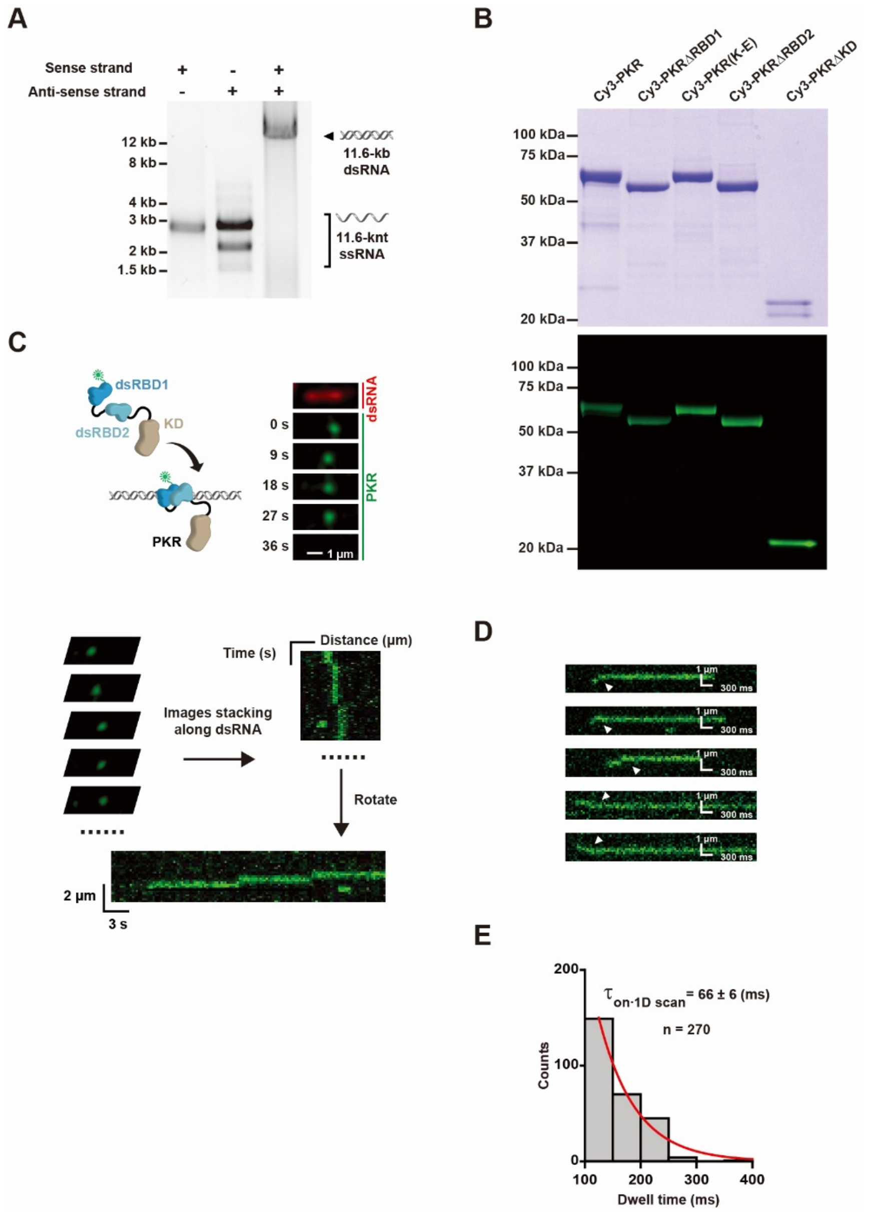


**Figure S1. The annealing of 11.6-kb dsRNA and fluorophore-labeled PKR.** (**A**) Agarose gel (0.75 %) showing the sense strand, anti-sense strand of ssRNA and the annealed 11.6-kb dsRNA. (**B**) Coomassie stained (top) and fluorescent images (bottom) of SDS-PAGE gels showing the purified PKR. (**C**) Top: A schematic illustration and representative fluorescent images showing the binding of a Cy3-PKR on 11.6-kb dsRNA. SYBR Gold stained 11.6-kb dsRNA is shown in red and Cy3-PKR at various times is shown in green. Bottom: An illustration of the kymograph construction. (**D**) Representative kymographs showing the “scan-and-trap” process at 50-ms imaging intervals. White arrowheads indicate the transitions from mobile binding to static binding. (**E**) Dwell time distribution of 1D scanning before PKR trapping (mean ± s.e., *n* = number of events). Data were fit to a single exponential decay to derive the average dwell time.

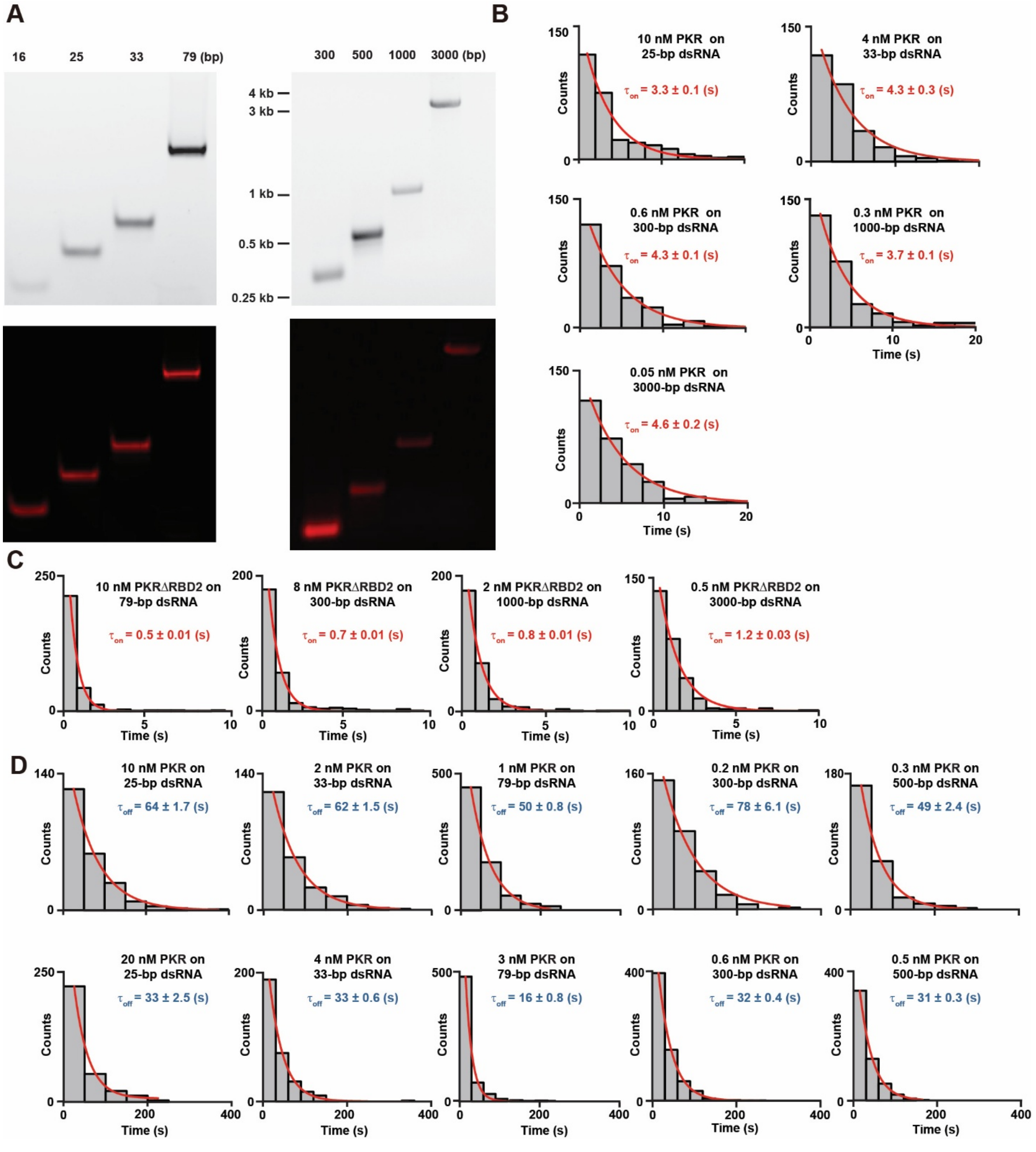


**Figure S2. Short dsRNA ligands used in co-localization assay, and the $\tau_{on}$ and $\tau_{off}$ of PKR binding to them.** (**A**) Left: Ethidium bromide stained (top) and fluorescent (bottom) PAGE showing 16-bp, 25-bp, 33-bp, and 79-bp dsRNA. Right: Ethidium bromide stained (top) and fluorescent (bottom) agarose gel showing 300-bp, 500-bp, 1000-bp, and 3000-bp dsRNA. (**B**) $\tau_{on}$ distribution of PKR on dsRNA of different lengths (mean ± s.e.). Data were fit to a single exponential decay to derive the average lifetime. (**C**) $\tau_{on}$ distribution of PKRΔRBD2 on various dsRNA ligands (mean ± s.e.). Data were fit to a single exponential decay to derive the average lifetime. (**D**) $\tau_{off}$ distribution of PKR on various dsRNA ligands (mean ± s.e.).

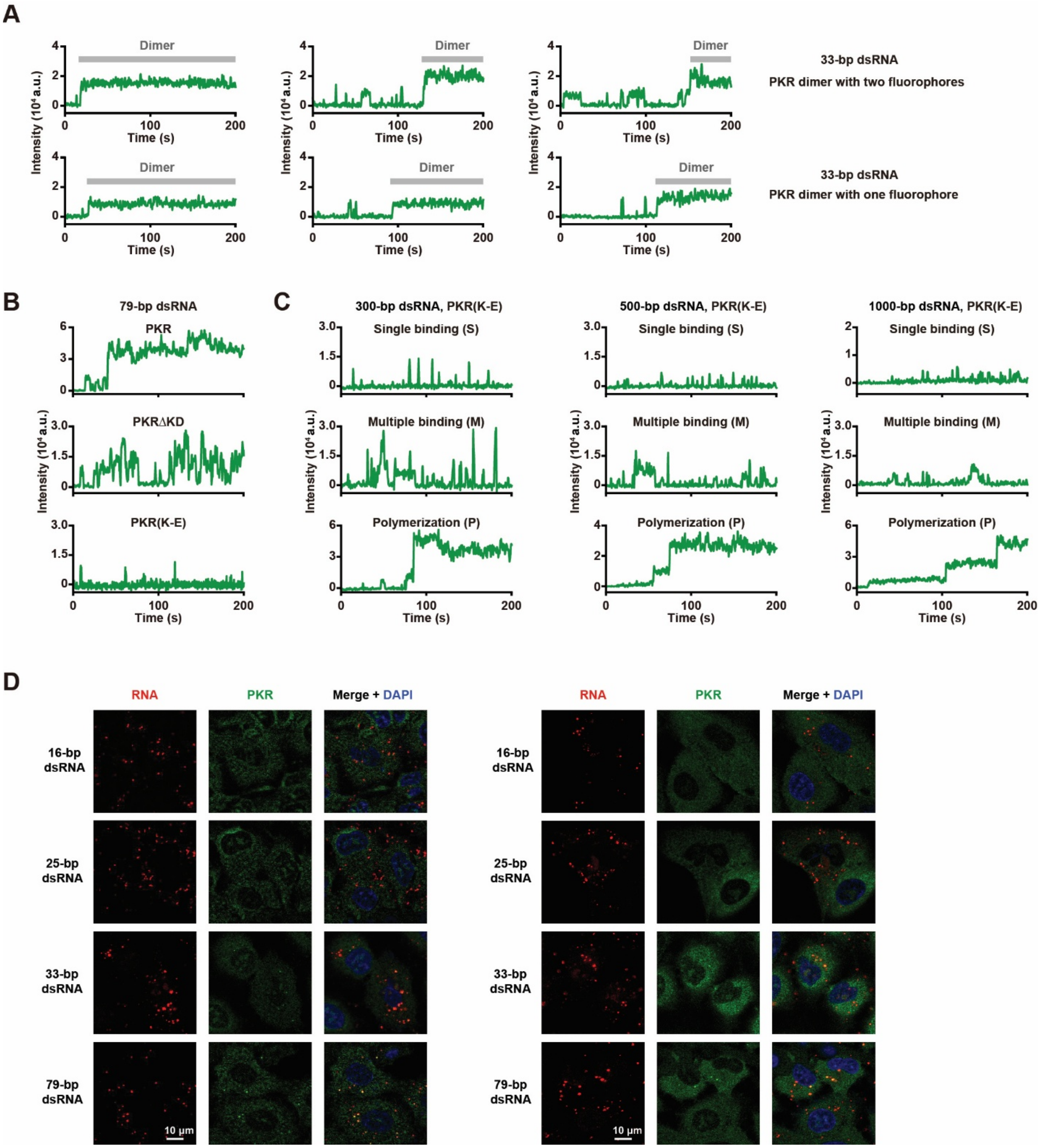


**Figure S3. Representative single-molecule trajectories and IF images.** (**A**) Representative trajectories showing the PKR dimers on 33-bp dsRNA with two (top) or one (bottom) Cy3 fluorophores. (**B**) Representative trajectories showing the PKR polymerization, PKRΔKD multiple binding and PKR(K-E) single binding events on 79-bp dsRNA. (**C**) Representative trajectories showing the single binding, multiple binding and polymerization events of PKR(K-E) on various dsRNAs. (**D**) Representative IF images showing endogenous PKR colocalized with 33-bp and 79-bp dsRNA, but not with 16-bp and 25-bp dsRNA in A549 cells.

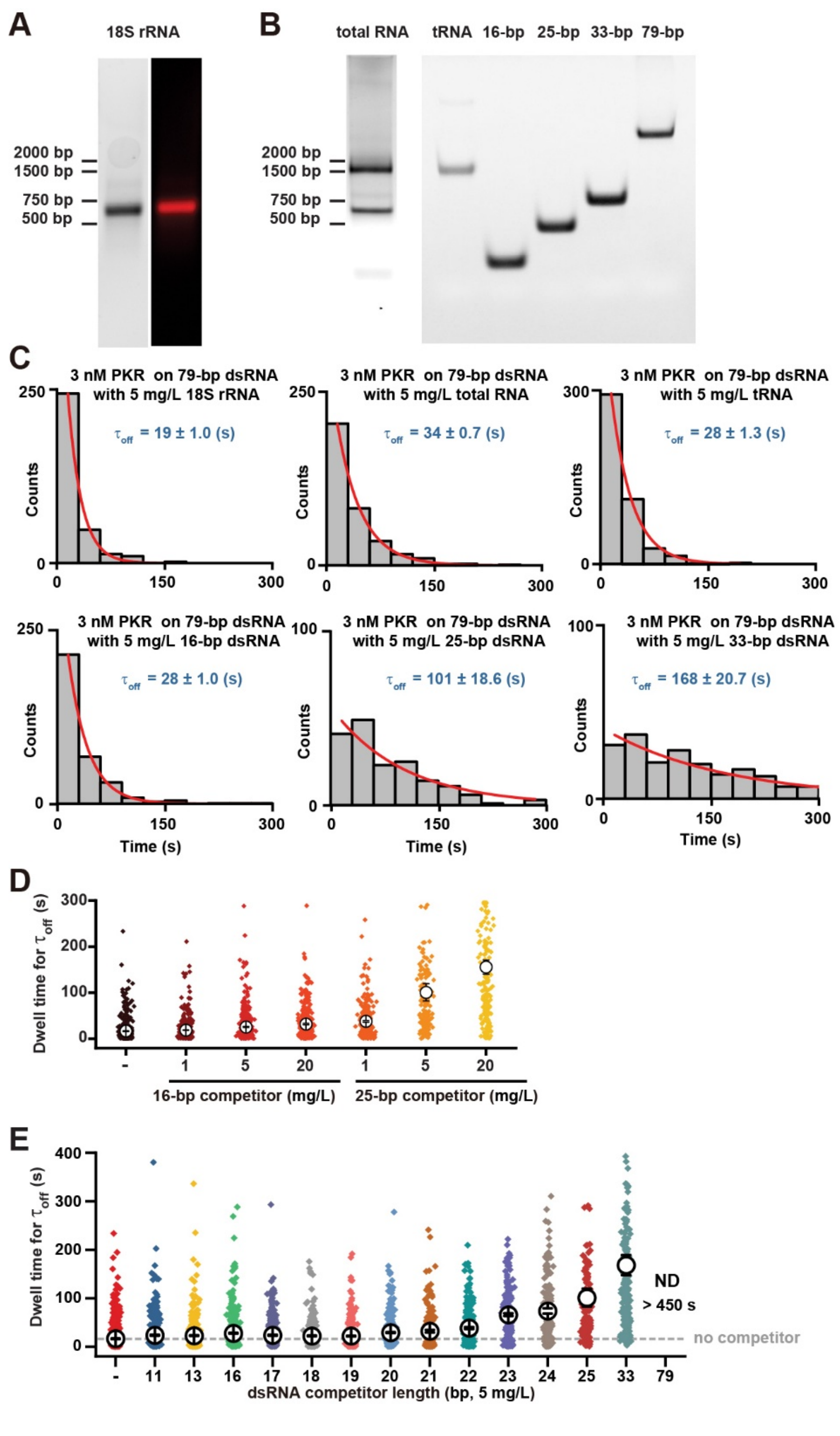


**Figure S4. 18S rRNA and RNA competitors used in co-localization assay, and the $\tau_{off}$ of PKR binding to 79-bp dsRNA in the presence of various competitors.** (**A**) Ethidium bromide stained (left) and fluorescent (right) agarose gel showing Cy5-labeled 18S rRNA. (**B**) Ethidium bromide stained agarose gel and PAGE showing total RNA, tRNA, 16-bp, 25-bp, 33-bp, and 79-bp dsRNA. (**C**) $\tau_{off}$ distribution of 3 nM PKR binding to 79-bp dsRNA in the presence of various

RNA competitors. Data were fit to a single exponential decay to derive the average lifetime (mean ± s.e.). (**D**) $\tau_{off}$ distribution of 3 nM PKR binding to 79-bp dsRNA with various concentrations of 16-bp or 25-bp dsRNA competitor. Diamonds represent individual events and open circles represent the mean dwell time derived by single exponential decay fitting (mean ± s.e.). (**E**) $\tau_{off}$ distribution of 3 nM PKR binding to 79-bp dsRNA with various 5 mg/L dsRNA competitors. Diamonds represent individual events and open circles represent the mean dwell time derived by single exponential decay fitting (mean ± s.e.).

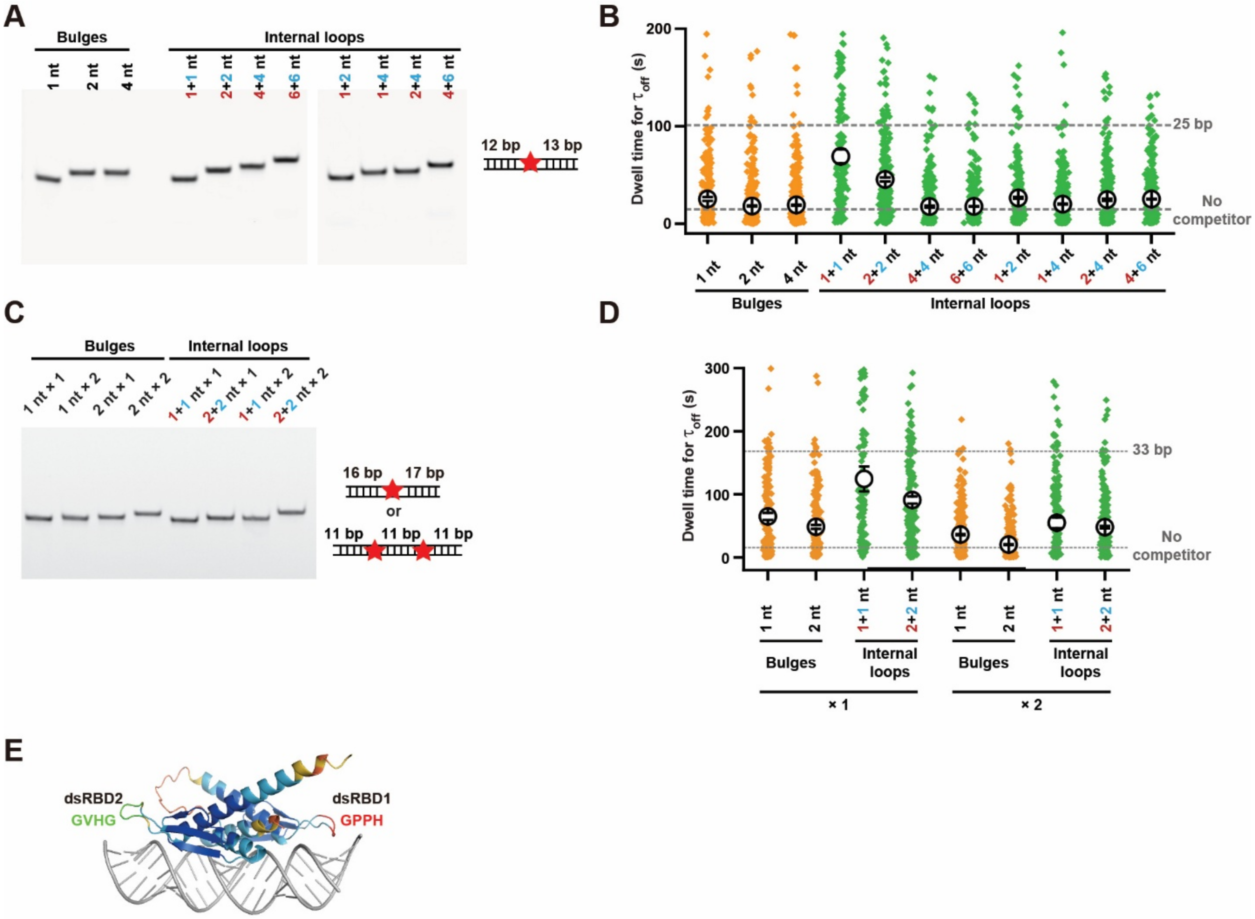


**Figure S5. Imperfect dsRNA ligands used in single-molecule studies, and the $\tau_{off}$ of PKR binding to 79-bp dsRNA with various imperfect dsRNAs.** (**A**) Ethidium bromide stained PAGE showing 25-bp dsRNA ligands with bulges or internal loops. (**B**) $\tau_{off}$ distribution of 3 nM PKR binding to 79-bp dsRNA with various imperfect 25-bp dsRNA ligands (5 mg/L). Diamonds represent individual events and open circles represent the mean dwell time derived by single exponential decay fitting (mean ± s.e.). (**C**) Ethidium bromide stained PAGE showing 33-bp dsRNA ligands with bulges or internal loops. (**D**) $\tau_{off}$ distribution of 3 nM PKR binding to 79-bp dsRNA with various imperfect 33-bp dsRNA ligands (5 mg/L). Diamonds represent individual events and open circles represent the mean dwell time derived by single exponential decay fitting (mean ± s.e.). (**E**) AlphaFold model of PKR's dsRBDs bound to a 25-bp dsRNA, showing that the canonical GPxH RNA-binding motif is present in dsRBD1 but absent in dsRBD2.
